# Direct Training of Networks of Morris-Lecar Neurons with Backprop

**DOI:** 10.1101/2025.11.23.690009

**Authors:** Navid Akbari, Kai Mason, Aaron Gruber, Wilten Nicola

**Affiliations:** Hotchkiss Brain Institute, University of Calgary, Calgary, Alberta, Canada; Cumming School of Medicine, University of Calgary, Calgary, Alberta, Canada; University of Calgary, Department of Cell Biology and Anatomy; Alberta Children’s Hospital Research Institute, University of Calgary, Calgary, Alberta, Canada; University of Calgary, Department of Physics and Astronomy

## Abstract

Spiking Neural Networks (SNNs) have the potential to replicate the brain’s computational efficacy by explicitly incorporating action potentials or “spikes”, which is not a feature of most artificial neural networks. However, training SNNs is difficult due to the non-differentiable nature of the most common spiking models: integrate-and-fire neurons. This study investigates if some of the difficulty in training SNNs arises from the use of integrate-and-fire neurons, rather than smoother alternatives, like conductance-based neurons. To that end, we considered networks of Morris-Lecar (ML) neurons, a conductance-based neuron model which is differentiable. Networks were built using kinetic synaptic models that smoothly link presynaptic voltage dynamics directly to postsynaptic conductance changes, ensuring that all components remain fully differentiable. Switching to biophysically detailed models of synapses and neurons enabled direct end-to-end training through Backpropagation Through Time (BPTT). Biophysically detailed networks were successfully trained on image classification, regression, and time series prediction tasks. These results demonstrate the feasibility of employing biophysically detailed differentiable point neuron models to create SNNs that function as more accurate paradigms for the study of neural computations and learning. Further, this work confirms that some aspects of the difficulty in translating gradient-based learning algorithms from machine learning may arise from model choice, rather than SNNs being intrinsically difficult to train.

**Author summary:** The brain’s information-processing efficiency arises in part from neurons communicating via discrete spikes. Spiking Neural Networks (SNNs) mimic this process at the neuronal level but have been difficult to train as most machine learning algorithms are not directly applicable. Most SNNs use integrate-and-fire neurons, a modelling framework that simplifies spikes into non-differentiable, abrupt voltage changes, which makes them difficult to train with powerful, standard AI training methods that use derivatives to compute gradients (e.g. Backprop). In our work, we asked if this difficulty could be overcome by considering end-to-end differentiable spiking neural networks. We used completely differentiable SNNs using the Morris-Lecar neuron, a biophysically detailed neuron model that produces smooth spikes, along with differentiable kinetic synapses. With the entire network being mathematically differentiable, we found that we could train it directly using standard backpropagation through time on different tasks (regression, classification, and chaotic time series prediction). This work demonstrates that the use of integrate-and-fire models may be limiting applications of machine learning algorithms towards understanding how learning functions in the brain.

## 2. Introduction

The human brain performs many of its operations and computations by using discrete events called action potentials or spikes [1]. Spiking Neural Networks (SNNs) are a family of Artificial Neural Networks (ANNs) that use spikes as an integral component in computation. The incorporation of an explicit mechanism of spiking within these networks allows flexible information encoding by multiple mechanisms [2, 3]. For example, neuronal firing rates, the relative timing of spikes, or sequences of spikes across networks are all encoding strategies that are observed in neural circuits both *in vivo* and *in silico* [4–6]

Despite their potential, training SNNs remains difficult [7]. Most SNN architectures employ simple neuron models, such as the Leaky Integrate-and-Fire (LIF) neuron, which features a discontinuous spike-generating mechanism at threshold [2, 8, 9]. LIF neurons have been used extensively in computational neuroscience as they have numerous advantages [10], such as the relative ease in mathematical analysis of networks [11–18], the efficiency of simulating large networks as models of neural circuits with millions of neurons [19, 20], neuromorphic applications where the hardware resources are limited [21]. However, integrate-and-fire neurons have one glaring deficiency which emerges in models of learning: they are not differentiable. The direct use of gradient-descent-based optimization techniques like Backpropagation Through Time (BPTT) [22], is made more difficult by this intrinsic non-differentiability. As a result, several different approaches to training SNNs have been devised that bypass this issue. These include event-based backpropagation techniques [23], methods that convert pre-trained ANNs to SNNs [19, 24], and the use of surrogate gradients to approximate the derivative of the spike function [9, 25, 26], or through the application of reservoir computing [8, 27, 28] using chaotic SNNs [29] as the computational basis. Despite the progress made in training integrate-and-fire based SNNs, existing learning algorithms often have to bypass the lack of differentiability in integrate-and-fire-based models.

In this work, we sought to determine if instead of bypassing the lack of differentiability in integrate-and-fire networks, we could instead utilize completely differentiable SNNs where gradient descent could be directly applied. This mirrors recent work on creating efficient gradient-based solvers for differentiable biophysical SNNs with the JAXLEY simulation framework [30]. Here, we considered an SNN architecture centered around the Morris-Lecar neuron model [31, 32], specifically leveraging its intrinsic differentiability, along with smooth synaptic connections utilizing kinetic synaptic models [33] which couple the presynaptic voltage to the post-synaptic conductance. The core motivation for selecting the network structure considered is its amenability to direct gradient calculation through end-to-end differentiability of the equations, thereby enabling the robust application of BPTT for network training. Further, the SNNs considered here are designed with biological plausibility at their forefront, notably incorporating separate populations of excitatory and inhibitory ML neurons within the hidden layer, consistent with Dale’s Law, where each neuron type establishes connections using its distinct synaptic effect. This approach, therefore, aims to bridge the gap between biologically plausible neuronal dynamics and effective gradient-based learning.

The proposed SNN architecture features a three-layer feedforward design, with the hidden layer containing balanced interconnected excitatory and inhibitory neurons that interact through conductance-based synapses to emulate biophysical synaptic transmission [33, 34]. Using completely smooth SNNs, we were able to successfully train SNNs to perform classification tasks (MNIST and Fashion-MNIST), nonlinear regression and the autonomous generation of different dynamical systems (Van der Pol Oscillator and Lorenz System). Through the successful training of these networks using direct BPTT, similar to the work in [30], we show that smooth SNNs not only function well on complex computational tasks but also provide a potentially more accurate representation of neural computation and learning in biological circuits while simultaneously promoting a closer relationship between computational neuroscience and machine learning. Our work demonstrates that at least some of the difficulty in training SNNs arises from the use of non-smooth integrate-and-fire neurons.

## 3. Results

### Comparison of Morris-Lecar and Leaky Integrate-and-Fire Neuron Dynamics

Many applications of machine-learning based algorithms have utilized leaky integrate-and-fire neurons, in some form or another [8, 9, 19, 25–28]. For one-dimensional integrate-and-fire neurons, the differential equation for the voltage of a neuron, *v*(*t*), is given by

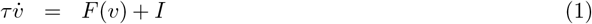

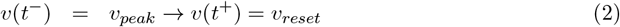

for some *F* (*v*), which determines the subthreshold dynamics of the voltage, *v*(*t*). In the case of the leaky integrate-and-fire neuron, *F* (*v*) = *−v* while other neuron models use nonlinear *F* (*v*) like the quadratic (*F* (*v*) = *v*^2^) or exponential (*F* (*v*) = *e*^*v*^*− v*) models. When *v*(*t*^*−*^) = *v*_*peak*_ at some time *t*^*−*^, the voltage is immediately reset (to *v*_*reset*_, Fig. 1-A). However, this produces a trade-off that emerges when we consider the powerful suite of gradient-based learning techniques that arise from machine learning: networks of LIF neurons are non-differentiable, and as a result, more difficult to train (Fig. 1-A). The non-differentiability also extends to synaptic models like the commonly used single exponential synaptic filter [13, 32, 33]:

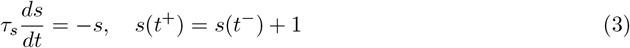

where the variables have discrete jumps (which are also non-smooth).

**Figure 1:**
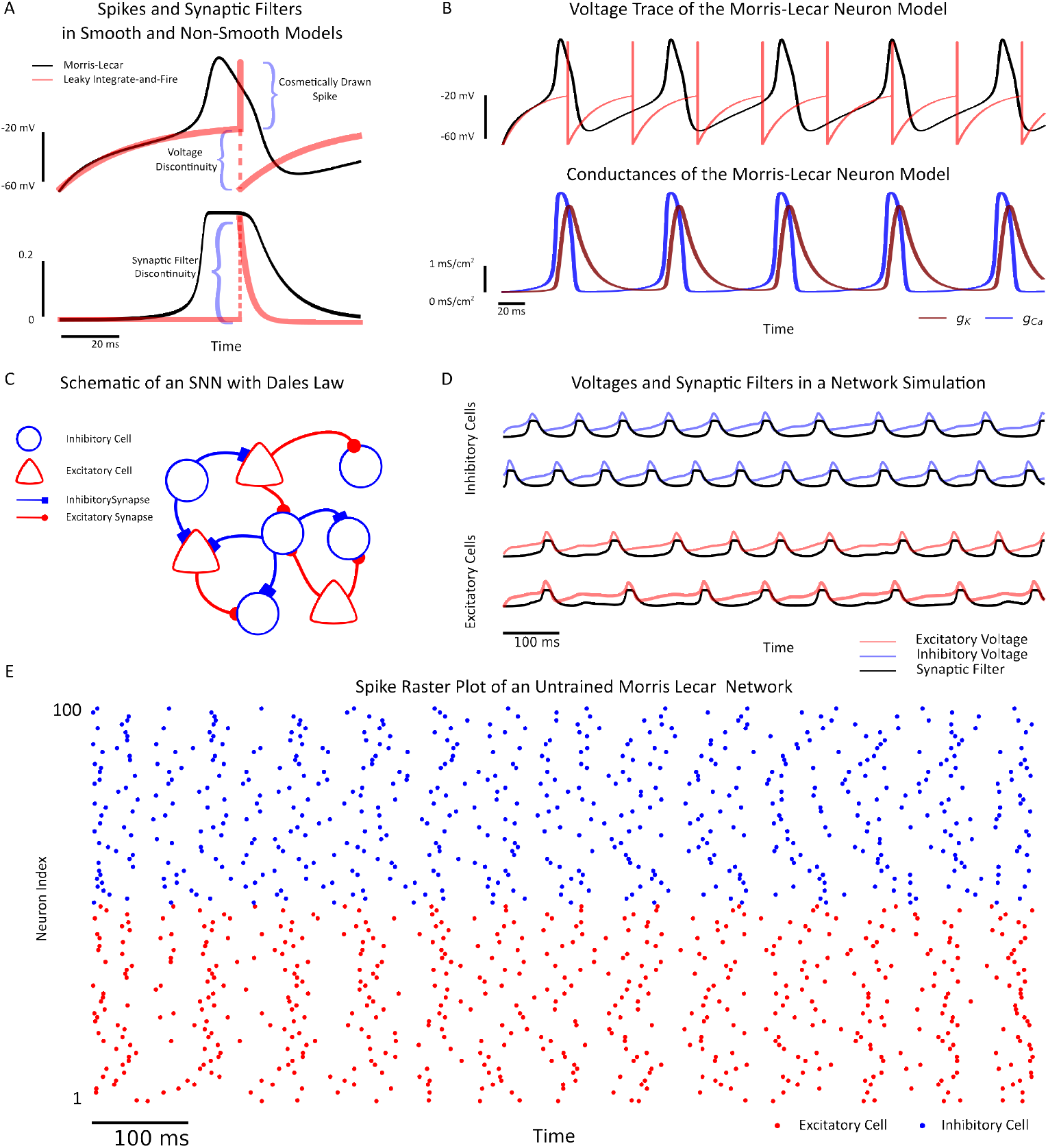
Morris Lecar Neurons and Networks are Differentiable. **A:** (Top) Single action potential shapes. The ML action potential waveform is smooth, whereas the LIF is discontinuous due to its instantaneous reset. The spike for the LIF model is cosmetically drawn. (Bottom) Synaptic filters. The single-exponential filter is discontinuous, while the gating variables of kinetic synapses are smooth functions of the presynaptic voltage. **B:** (Top) Representative voltage traces, illustrating subthreshold and smooth spike dynamics in the ML model compared to the simpler LIF threshold-reset behavior. (Bottom) Dynamics of ion conductances (*g*_*Ca*_(*t*), *g*_*K*_(*t*)) responsible for the ML neuron’s smooth voltage trajectory. **C:** Schematic of a small randomly configured excitatory/inhibitory network with conductance-based synapses. **D:** Voltage traces (coloured) and synaptic filter dynamics (black) in the ML network (excitatory/red and inhibitory/blue) simulation with a balanced network consisting of *N* = 100 neurons. **E:** ML network raster plot.

To investigate if integrate-and-fire models are the source of some of the consternation in applying gradient-based learning to spiking neural networks, we considered a class of smooth and differential spiking models: conductance-based neuron models. This class of models was developed from the classical Hodgkin-Huxley model [1] to capture the underlying biophysics of action potential generation and transmission. Here, we specifically focus on the Morris-Lecar neuron model:

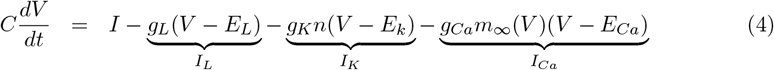

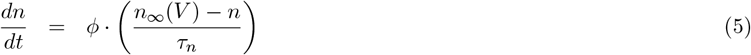

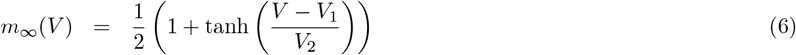

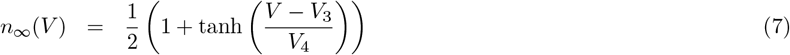

due to its differentiable nature and efficient-to- simulate two-dimensional dynamics. The voltage for the ML neuron is given by *V* (*t*) while *n*(*t*) acts as the gating variable for the potassium gate, which smoothly resets spikes. The ML neuron also features a series of conductances for Potassium (*g*_*k*_), Calcium 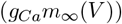, and a general leak-conductance (*g*_*L*_) that control the amount of current flowing through a neuron, while the reversal potentials (*E*_*k*_, *E*_*Ca*_, *E*_*L*_) in conjunction with the voltage dictate the direction of current flow.

At the single-neuron level, Fig. 1-A shows single spike shapes, with the ML model spike exhibiting a differentiable and (therefore) continuous spike. In contrast, the LIF model spike displays an instantaneous reset after reaching the threshold, which produces a sharp and discontinuous spike [35]. This discontinuous spike poses a challenge for gradient-based learning methods like back-propagation through time (BPTT) due to its non-differentiability at the spike threshold. In contrast, the ML neuron model’s smooth dynamics offer more biological plausibility and can be used in gradient-based learning methods, as illustrated in its voltage trace in Fig. 1-A&B. The ML neuron model’s differentiability comes from the underlying conductances (Fig. 1-B) implementing the spike generation/reset mechanism [36, 37].. The primary trade-off in using conductance neurons is that the additional variables and nonlinear operations required to simulate spikes biophysically implies that they are more computationally intensive to simulate.

To analyze network-level phenomena, we connected model neurons with conductance-based or ‘kinetic’ synaptic coupling [33, 34, 37] (Fig. 1-C&D):

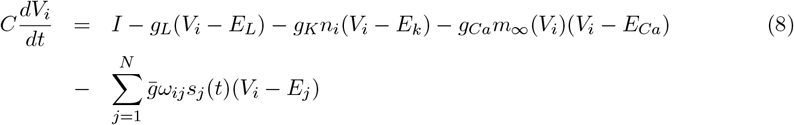

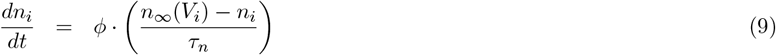

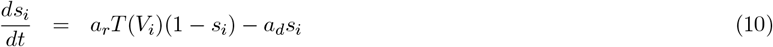

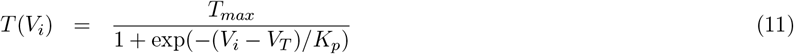

where 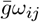 acts as the unitary synaptic conductance connecting neuron *j* to neuron *i*, and *E*_*j*_ is the reversal potential for the synapses formed by neuron *j*. The networks considered had balanced inhibitory/excitatory population of neurons; each neuron type had connections to all neurons (including autapses), with random connection strengths. The unitary synaptic conductances were generated from a uniform distribution in the interval 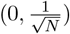 where *N* was the total population size although other potential scalings on the unitary conductance are possible [38], along with other initial configurations of the adjacency matrix [39]. The coupled ML neural network displayed irregular spiking dynamics Fig. 1-E in the initial coupling state (Supplementary Figure 1). Note that equations (8)-(11) are also differentiable for all of their parameters.

### Gradient Descent

In order to use conductance-based SNNs for learning, a readout from the network first needs to be defined:

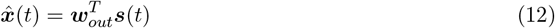

where 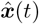 acts as the approximation to some *k* dimensional target, ***x***(*t*), and ***w***_*out*_ is some *N* × *k* dimensional matrix that projects onto the synaptic gating variables ***s***(*t*). Note that a constant bias term was also incorporated into the readout weights (Materials and Methods).

Next, we define the loss as:

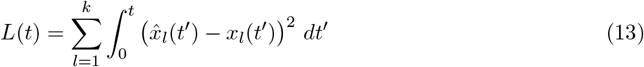

for continuous time tasks, or as

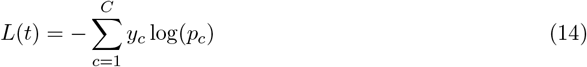

for classification tasks. The latter loss is conventionally called the Cross-Entropy Loss [40]. In this loss function, *y*_*c*_ represents the ground-truth label and is a standard one-hot vector:

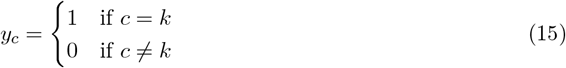

Where *k* is the index of the correct class. The term *p*_*c*_ is the model’s predicted probability for a given class *c* and is computed from the time-averaged network readout. First, we define the average readout (or logit) 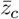 for each class *c* over time *t*:

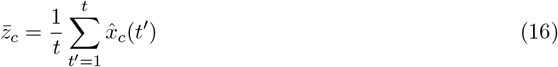

where 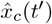 is the network readout for class *c* at time *t*^*′*^. The probability *p*_*c*_ is then calculated by applying the softmax function to these average logits:

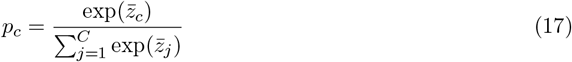

With the readouts and loss both defined, gradient-based training in SNNs can be accomplished by determining the partial derivatives of the loss as a function of the network parameters, and then iteratively updating the parameters with these gradients. The parameters utilized in the learning were the readout weights (***w***_*out*_), the recurrent weights (***w***) and the input weights ***w***_*in*_. In addition, all neurons received bias currents which were also learned.

Owing to the end-to-end differentiability of this system, the partial derivatives of the loss with respect to the free parameters 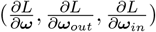 were readily derived using Backpropagation Through Time (BPTT) (see Supplementary Derivation). However, the actual weight updates were conducted with the AdamW optimizer and Pytorch approximating the partial derivatives for computational efficiency of the implementation (Materials and Methods). One of the advantages of considering smooth synapses and neuron models is that existing numerical optimizers can be considered owing to the end-to-end differentiability of these networks [30]. With the gradients in hand, all parameters are updated with:

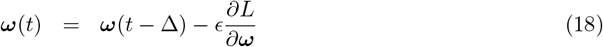

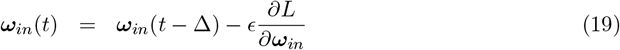

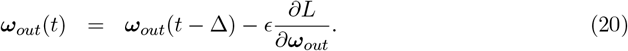

Note that full BPTT can be resource-intensive, especially for long temporal sequences, as it needs to store the gradients for every time step in the forward and backward passes. To mitigate these computational demands, we used Truncated Backpropagation Through Time (TBPTT) [41].

### Image Classification in Networks of Smooth Spiking Neurons

With the network and gradient defined, we evaluated the performance of the smooth Morris-Lecar SNN on standard machine learning tasks. We used a feedforward architecture between layers and incorporated coupled ML neurons in the hidden layer, evaluating performance on two widely used image-classification benchmarks: MNIST and Fashion-MNIST. These intra-layer synaptic connections created recurrent dynamics, allowing the neurons within the hidden layer to influence each other over time (Figure 2). For these tasks, the input to the network at each discrete time step of the simulation consisted of a vector of normalized pixel intensities of the images. Presenting the raw spatial input over a fixed duration T enabled the network to process the data continuously, allowing the coupled hidden-layer neurons to interact dynamically and extract spatial features needed for pattern recognition. A simple 10-dimensional readout matrix projected onto the hidden layer to extract the network output. The supervisory signal that was utilized to train the network consisted of a constant value of 1 for the output neuron corresponding to the correct class, while other output neurons remained inactive, set to a constant value of 0.

**Figure 2:**
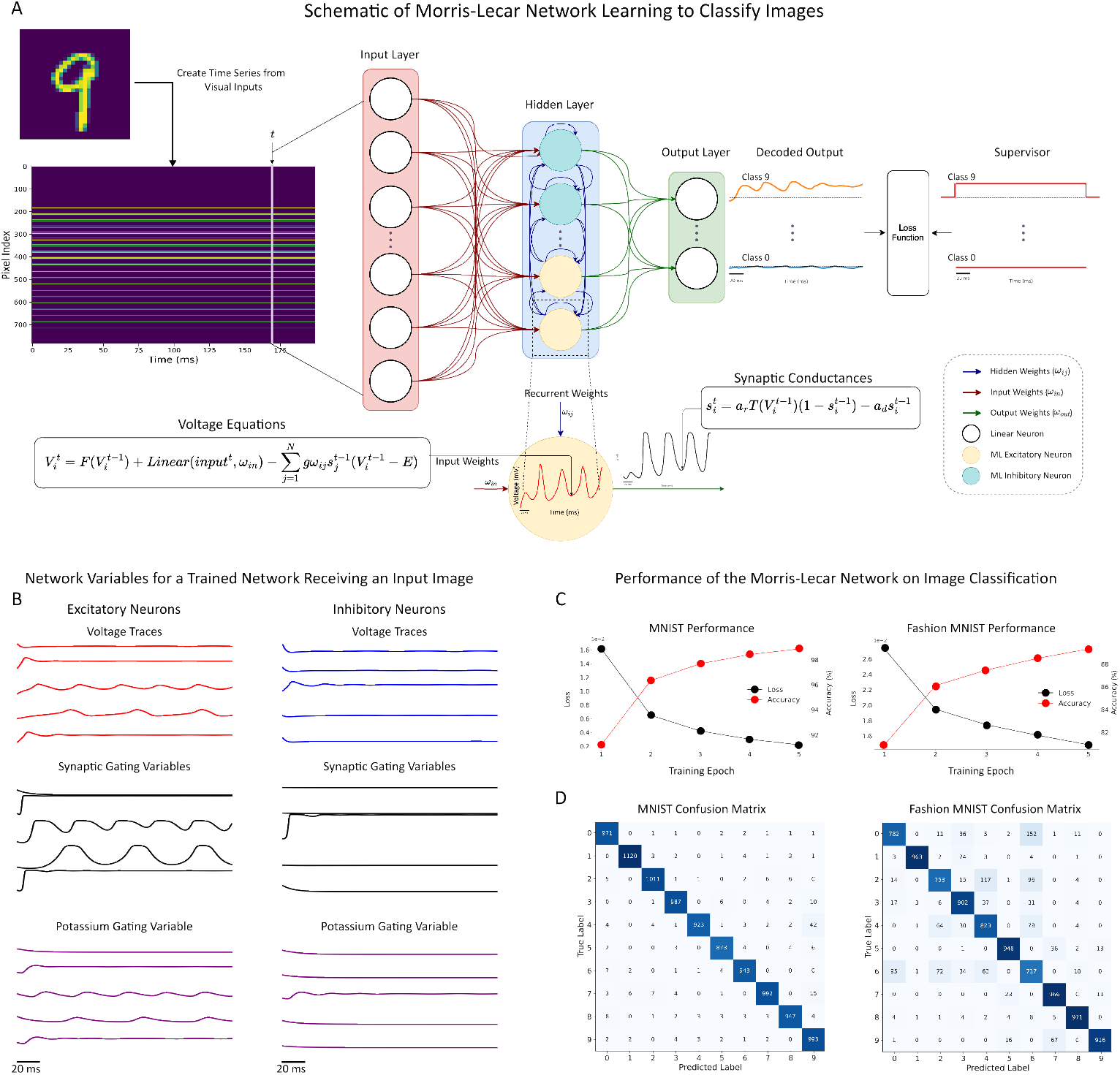
Performance of the ML-SNN on MNIST and Fashion-MNIST Classification. **A:** Schematic of a Morris-Lecar network receiving input images and classifying these images into discrete classes (in this case for the MNIST digit recognition task). **B:** Network variables upon receiving an input for inhibitory (left) and excitatory (right) neurons. The variables include the voltage (top), the synaptic gating variables (middle), and the Potassium gating variable (bottom). **C:** Training loss (black) and accuracy (red) curves over 5 epochs for the MNIST dataset. Training loss (black) and accuracy (red) curves over 5 epochs for the Fashion-MNIST dataset. **D:** (Left) Confusion matrix for the MNIST test set, achieving 97.60% accuracy (Right) Confusion matrix for the Fashion-MNIST test set, achieving 87.41% accuracy.

The network architecture for both of the classification tasks consisted of an input layer with 784 neurons, as both datasets’ image size was 28 × 28. The network also used a hidden layer with 1000 ML neurons with 500 interneurons and 500 pyramidal neurons, and an output layer with 10 readout neurons. The training was conducted for 5 epochs in both data sets.

We first evaluated the performance of the ML-based SNN on the well-known MNIST dataset [42], which consisted of images of handwritten digits from 0 to 9. This dataset is a widely used benchmark in the machine learning literature for classification tasks in model development. This dataset was structured with 60, 000 training images and a separate set of 10, 000 images for testing the performance of the model.

We used the training dataset to train the model using TBPTT to update the model’s synaptic weights and other trainable parameters. As the training phase shows, Fig. 2 A-C, in the first epoch, the model achieved an accuracy of 91.20%. At the end of the last epoch, the model’s accuracy increased to 98.88%.

For the test set, the model achieved 97.60% accuracy. As shown in Fig. 2-D, the confusion matrix details the model’s prediction versus the true labels. The high accuracy of the model is reflected in the elements on the diagonal, which indicate the number of correctly classified images. This accuracy is consistent with the performance of a single-layered artificial neural network with a comparable number of (artificial) neurons [43].

To further evaluate direct gradient descent in smooth SNNs, we trained the ML network on the Fashion-MNIST dataset [44]. Fashion-MNIST contains simple images of clothing items that need to be categorized into discrete classes. Similar to the MNIST dataset, this dataset comprised a 60,000-image training set and 10,000 testing images.

The same model architecture was used for this task, and we trained the model for 5 epochs. As illustrated in Fig. 2-C, final accuracy on the training set was 89.34%. Using this trained model, we evaluated the test dataset, which resulted in 87.41% accuracy. The confusion matrix, Fig. 2-D, shows a detailed breakdown of the model’s predictions versus the true labels. This level of accuracy is also consistent with ANN/MLP based classifiers applied to the fashion-MNIST dataset [44].

Overall, the Morris-Lecar network trained with gradient descent could readily learn image classification on both the MNIST and Fashion-MNIST datasets by directly applying backpropagation through time.

### Learning Nonlinear Transformations in Networks of Smooth Spiking Neurons

Next, we investigated if the end-to-end differentiability of the smooth SNNs considered would allow for learning in a simple non-linear regression task. For this task, the model was trained to approximate the quadratic function *f* (*x*) = *x*^2^ for inputs *x ∈* [*−*2.0, 2.0]. We generated a dataset consisting of 5000 inputs, 80% of which used for training the model and the rest for testing. The network architecture was similar to the classification model using *N* = 1000 neurons (500 *E* and 500 *I*), but with a single input neuron and a single linear readout neuron. During training and inference, a constant input value *x* was presented to the network for a duration of *T* = 500 ms.

The network successfully learned the target nonlinear mapping, as shown in Fig. 3-A. The average mean squared error (MSE) loss decreased from 66.83 at first to 5.80 by epoch 20 while the Mean Absolute Error (MAE) similarly improved from 0.94 to 0.10 during learning indicating the convergence to the target function. The network’s final output for a given input *x* was computed by averaging the network’s output signal over the latter 250 ms of the simulation window, as illustrated in Fig. 3-B. This time-averaging process translated the continuous spiking dynamics into a single-value estimate while removing transients that occurred when new inputs were provided to the network. The internal network dynamics underlying this computation are shown in Fig.3-C, which shows that the hidden layer neurons settle into a stable pattern of activity for each input.

**Figure 3:**
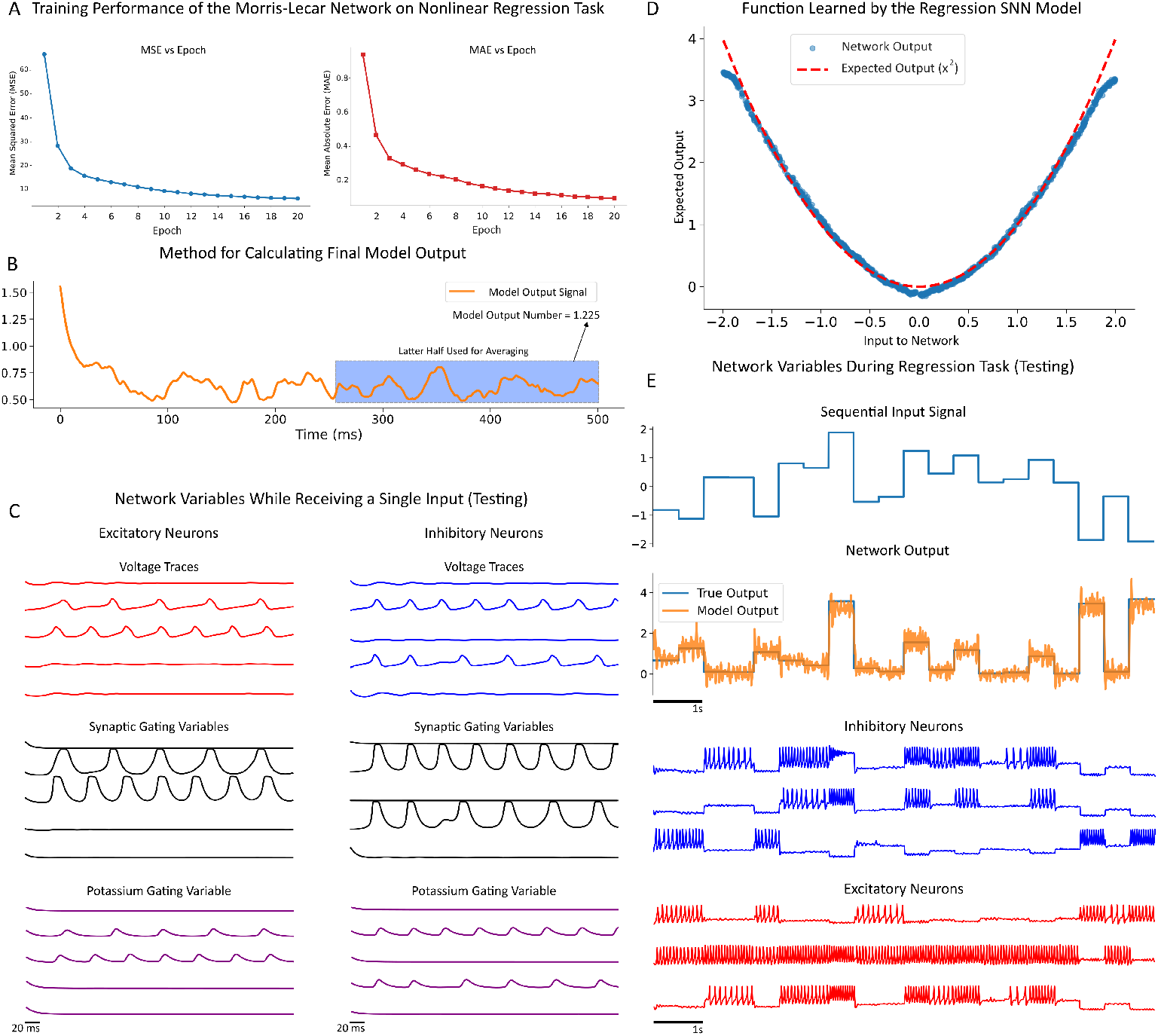
ML-SNN Performance on Static Function Approximation Task. **A:** Training performance over 20 epochs, showing a rapid decrease in the time-distributed Mean Squared Error (MSE) loss (left) and the Mean Absolute Error (MAE) of the final number prediction (right). **B:** Method for calculating the model’s number output. The instantaneous model output signal (orange) fluctuates over the 500 ms simulation. The final prediction (model output number) is calculated by averaging the signal over the latter 250 ms (blue shaded region) **C:** Internal network variables for a single input. Traces from representative excitatory (left, red) and inhibitory (right, blue) neurons, showing (from top to bottom): Voltage, synaptic gating variable, and potassium gating variable; showing that network settles into a stable dynamic state. **D:** Generalization performance on the test set. The network’s model predictions (blue dots) are plotted against the true function (red dashed line), demonstrating a highly accurate approximation of the parabola. **E:** Network response to a piecewise-constant sequential input signal (top). The instantaneous model output (middle, orange) closely tracks the true output (middle, teal). The corresponding voltage traces for inhibitory (blue) and excitatory (red) populations (bottom) show how firing patterns modulate in real-time to represent the changing input.

A clear validation of the model’s generalization on the test set is shown in Fig. 3-D, showing the network’s predictions (blue dots) against the true values of the target function (red dashed line). Quantitatively, the model achieved an MAE of 0.09 and an average MSE loss of 5.73 on the test set, consistent with the training loss.

During testing, we presented a sequential series of inputs from the test set (Fig. 3-E top), each lasting approximately 500 ms. The network’s output (Fig. 3-E middle) effectively tracked the target *x*^2^ values, rapidly adjusting its mean activity level as the input changed. The corresponding voltage traces (Fig. 3-E bottom) show how the firing patterns of both excitatory and inhibitory populations modulate in real-time to represent the different input levels. This result confirms smooth SNNs capacity not only for pattern recognition (Fig. 2) but also for precise nonlinear function approximation.

### Learning Autonomous Dynamics in Networks of Smooth Spiking Neurons

In addition to the pattern recognition and regression tasks, we also considered direct application of TBPTT to train smooth Morris-Lecar networks to generate complex temporal dynamics and patterns autonomously. By slightly modifying the model architecture and feeding back 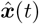 to the neurons in the hidden layer, the network formed a closed-loop model that can generate learned dynamics only by using its own internal states and output. We tested the ML-SNN’s ability to learn and replicate the behavior of two well-known dynamical systems: the Van der Pol oscillator and the chaotic Lorenz system. These tasks evaluated the capacity for ML-SNNs to generate time-dependent sequences by leveraging the rich internal dynamics of the ML hidden-layer neurons.

We employed a supervised learning technique during the training phase, in which the network was trained to predict the state variables of the system in the next time step. Using the Mean Squared Error (MSE) loss function, the network’s output was compared to the supervisory signal to compute the gradient in TBPTT.

For the first task of dynamical system reconstruction, we used the Van der Pol oscillator, which is characterized by stable limit-cycle oscillation with two variables and a parameter (*µ* = 1):

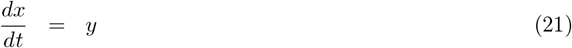

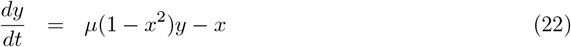

The network consisted of 500 neurons with 250 excitatory and 250 inhibitory Morris-Lecar neurons. After training, we tested the performance of the SNN using identical initial values for the model’s 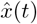 and the true Van der Pol system at time *t* = 0. The SNN successfully generated the dynamics solely based on its own internal state variables and learned parameters, with the output at each time step serving as the input for the next. As illustrated in Fig. 4A-B, the network successfully generated the Van der Pol dynamics autonomously and captured the characteristic oscillatory behavior.

**Figure 4:**
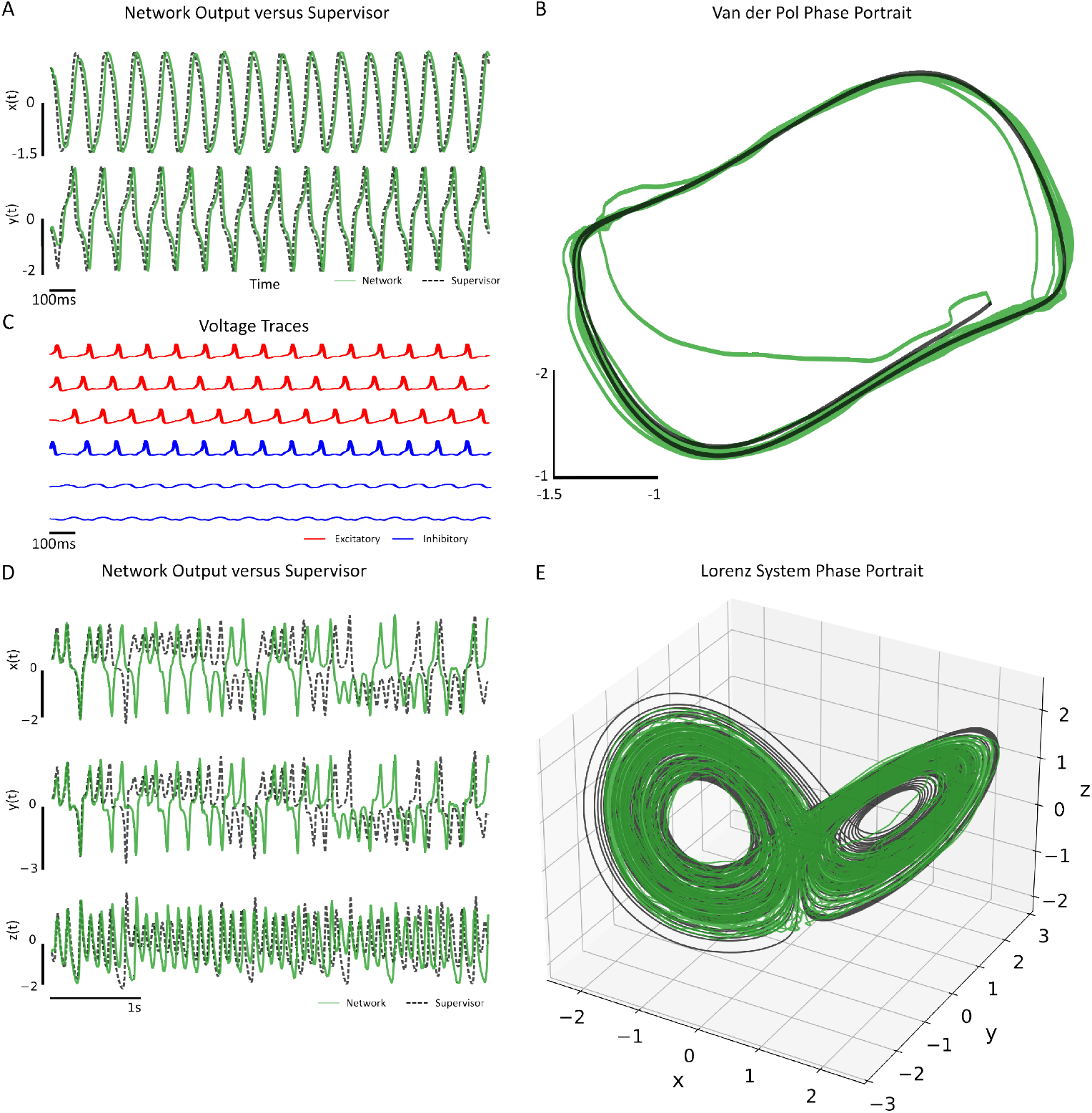
Autonomous Generation of Van der Pol and Lorenz Dynamics using Conductance Based SNNs trained with BPTT A-C: Van der Pol oscillator Results. (**A**) Time series of the *x* and *y* variables, comparing true (black solid) and generated (green dashed) trajectories. (**B**) Phase plane (*x* vs *y*) comparison of true (black) and generated (green) limit cycles, showing close overlap. (**C**) Random Neurons voltage traces during autonomous generation of Van der Pol dynamics **D-E:** Lorenz system results. (**D**) Time series of the *x, y*, and *z* variables, comparing true (black dashed) and generated (green solid) trajectories. (**E**) 3D phase space comparison showing the generated trajectory (green) recreating the geometry of the true Lorenz attractor (black).

As shown in Fig. 4-A, a visual analysis demonstrated a strong correlation between the autonomously produced state variable trajectories by the SNN (dashed lines) and the true trajectories (solid lines), signifying precise synchronization in both phase and amplitude. Importantly, the phase plane depiction in Fig.4-B validates that the network effectively replicated the characteristic limit cycle attractor of the Van der Pol system, with the generated path (green) closely mirroring the actual path (black). This finding underscored the capability of the ML-based SNN, when suitably trained, to acquire and consistently maintain nonlinear oscillatory dynamics.

Next, we considered a more difficult dynamical systems task, chaotic time-series prediction. We assessed the network’s ability to emulate the chaotic Lorenz system (Figure 4-D&E), characterized by its three parameters 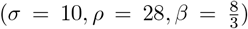 and governed by the differential equations:

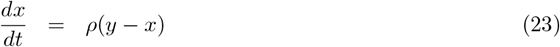

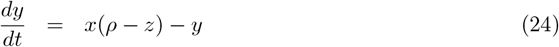

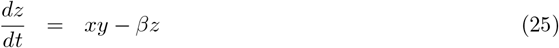

As before, the network was trained with the AdamW optimizer and the gradients were numerically estimated with TBPTT.

After the training phase, we evaluated the autonomous generation capability of the model. We fed an initial condition to the network and allowed the network to operate in a closed-loop mode for 20 s, as shown in Fig. 3-D for 5 s for better illustration. The time series produced Lorenz-like trajectories for the trained network, while the attractor of the network was a noisier version of the Lorenz butterfly attractor, indicating successful training. The voltage traces of individual neurons were also encoding the system states in both spiking and sub-threshold modes (Supplementary Figure 2). Finally, we remark that the return map, a plot of the 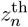 peak against the 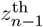 peak was also reproduced by the Morris-Lecar network with noise induced by spiking (Supplementary Figure 3).

Collectively, these results show that Morris-Lecar based SNNs with differentiable synapses and spike generation mechanisms can be trained as if they were classical artificial neural or recurrent neural networks with direct application of existing tools (i.e. PyTorch/AdamW) for estimating gradients. Some of the difficulty in training SNNs comes from the use of integrate-and-fire neurons, explicitly, rather than spiking itself.

## Discussion

Learning in spiking neural networks still remains challenging, despite recent advances. This study considers whether some of the difficulties arise from using integrate-and-fire models in understanding gradient-based learning in the brain, rather than anything intrinsic to spiking itself. We showed that it was possible to train biophysically detailed neural networks consisting of coupled excitatory/inhibitory neurons with smooth conductance-based synapses and a smooth spike generation mechanism. These networks were trained on image classification, regression, and dynamical systems tasks successfully. The training was with the conventional Backpropagation Through Time algorithm [22] that was directly applied. This was enabled by using direct gradient calculation via automatic differentiation. Our work complements recent simultaneous efforts with the JAXLEY framework that focus on training and differentiating morphologically detailed biophysical models by using direct gradient descent to fit to neural data [30]. These models were also trained with direct automatic differentiation and coupled with conductance based synapses. Collectively, our results and [30] demonstrate the feasibility of training biophysically detailed networks directly with backpropagation. We found that the primary tradeoff in using a Morris-Lecar model was that the conductance-based networks took longer to simulate and train, largely due to the increased computational complexity of modelling spikes explicitly. Although the ML model requires more floating operations per second (FLOPs) than the LIF model, our choice of this model was deliberate and central to our focus on biological plausibility [20]. The ML neuron is a conductance-based neuron which models the dynamics of voltage gated ion channels as spikes are generated [31]. These mechanistic dynamics enable the ML model to exhibit different neuronal behaviours, including both Type II and Type I excitability, which is characterized by unique phase-responses and network synchronization effects in contrast to the LIF model [32, 45, 46].

Training LIF-based SNNs requires computational methods to bypass or circumvent their non-differentiability. For instance, ANN-to-SNN conversions are powerful methods for constructing functional spiking networks [19, 24, 47, 48] but seldom consider spike times [3, 49, 50]. Surrogate gradient methods, which replace the non-smooth functions that emerge in network equations due to spike discontinuities with smoother differentiable functions have been applied in direct SNN training for both task domains [9, 25, 51, 52] and can also use spike times explicitly for coding [53]. Event-based backpropagation is used for closer alignment with sparse operation in pattern recognition [23]. Reservoir based methods attempt to bypass full gradient calculation altogether [8, 27, 28], and typically utilize recursive-least-squares variants as a kind of feedback controller to stabilize network dynamics onto a target system. We highlight that these methods have emerged as the computational benefits of using integrate-and-fire models are considerable. Indeed, very large LIF networks can be simulated to overcome any potential accuracy issues, and the LIF neuron serves as a kind of “neuromorphic primitive” that is easily synthesized in hardware.

The approach we consider here is better suited for investigations of learning/gradient descent in populations of neurons, rather than applications of SNNs in neuromorphic hardware or large-scale simulations. Indeed, a central limitation of this work is the amount of computing resources required to simulate networks of conductance neurons with conductance-based synapses in comparison to LIF networks. Non-smooth models require significantly less FLOPS (floating-point operations per second) than even the Morris-Lecar neuron, one of the simplest conductance based neurons [20, 54]. This implies that while using conductance-based neurons with conductance-based synapses will allow us to test the hypotheses that biological neurons may use different gradient-based learning algorithms [55, 56], these types of networks will be challenging to synthesize in either digital or analog neuromorphic implementations due to the computational resources required. However, we do remark that large-scale simulations of biophysical detailed (and parameter constrained) neurons as in [30] or the point neurons considered here are certainly feasible with high-performance computing.

Although the ML model has been integrated into SNNs in prior research, including investigations into STDP-based unsupervised learning within memristive networks [57] and training using the differential evolution algorithm [58], these methodologies focused on distinct learning paradigms and goals. Here, we computationally demonstrate that conductance based Morris-Lecar networks coupled with conductance based synapses can be readily trained on a variety of fundamental tasks with stock BPTT.

## Acknowledgments

We thank Ben Livingstone and Jafar Shamsi Chamyousefali for their contributions.

## Code Availability

ML-SNN implementation for all the tasks can be found here: https://github.com/navidakbari/ML-SNN

## Materials and methods

In this section, we detail the Morris-Lecar (ML) neuronal model, conductance-based synaptic coupling, and the overall architecture of the SNN model.

### Morris-Lecar Neuron Model

The Morris-Lecar neuron model is a two-dimensional neural model that exhibits differentiability. The ML model is driven by three essential ionic currents: membrane leakage (*I*_*L*_), potassium (*I*_*K*_), and calcium (*I*_*Ca*_) currents. The following equations show the simplified ML model:

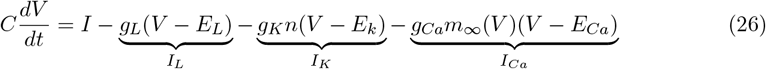

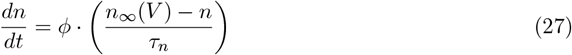

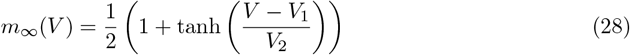

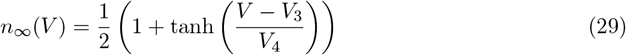

Variables *V, I*, and *n* in Eq. 26 & 27 represent the membrane potential, total membrane current, and the fraction of an open potassium ion channel. Furthermore, *m*_*∞*_ and *n*_*∞*_ are the calcium and potassium ion channels conductance at their steady-state given *V*, respectively. Finally, the term *τ*_*n*_ corresponds to the rate constant governing the opening of the potassium channel. Note that conventionally *τ*_*n*_ is voltage dependent as well. However, we fixed it due to numerical overflow errors when *τ*_*n*_ is voltage-dependent [35]. The model’s parameters and their values are described in Table 1. Using the ML neuron model described above, we initialized the SNNs considered as a balanced excitatory/inhibitory neuronal population network.

**Table 1:** Morris-Lecar neuron parameter values and description.

|  |  |
| --- | --- |
| $C = 20.0$ ($\mu\text{F}/\text{cm}^2$) | Membrane capacitance |
| $g_L = 2.0$ ($\text{mS}/\text{cm}^2$) | Leak conductance |
| $g_K = 8.0$ ($\text{mS}/\text{cm}^2$) | Maximum conductance of potassium channels |
| $g_{Ca} = 4.0$ ($\text{mS}/\text{cm}^2$) | Maximum conductance of calcium channels |
| $E_L = -60.0$ (mV) | Leak reversal potential |
| $E_K = -84.0$ (mV) | Potassium reversal potential |
| $E_{Ca} = 120.0$ (mV) | Calcium reversal potential |
| $\phi = 0.067$ | Rate scaling constant |
| $V_1 = -1.2$ (mV) | Scaling parameter |
| $V_2 = 18.0$ (mV) | Scaling parameter |
| $V_3 = 12.0$ (mV) | Scaling parameter |
| $V_4 = 17.4$ (mV) | Scaling parameter |

Morris-Lecar neuron model parameters values based on the class I spiking regime [59]. Units are provided in parentheses where applicable.

### Connecting ML Neurons into Networks

To establish connections among Morris-Lecar neurons, we utilized a conductance-based synaptic coupling [32, 33] where the presynaptic voltage directly and smoothly impacts the postsynaptic conductance of a synapse.

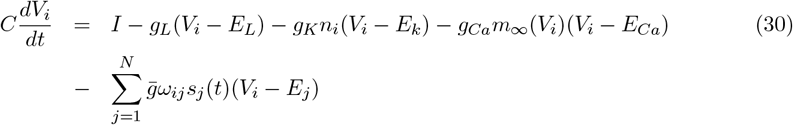

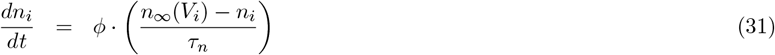

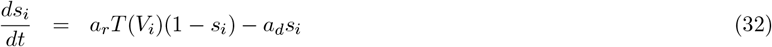

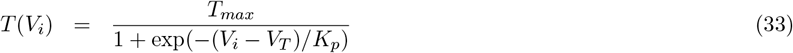

The total membrane current and ionic currents in Eq. 26 were combined into *F* (*V*_*i*_). These ML neurons were interconnected using a smooth synaptic gating variable described in Eq. **??**. The synaptic connection from neuron *j* to neuron *i* was represented as 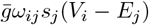, where the synapse’s inhibitory/excitatory nature was determined by the reversal potential *E*_*j*_, and the network parameter values are displayed in Table 2. The neurotransmitter release from neuron *I* into the synaptic cleft was modeled by the function *T* (*V*_*i*_), detailed in Eq. **??**. The conductance matrix *ω* was an *N* × *N* matrix where all unitary conductances were positive with the following constraint:

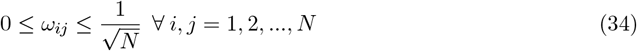

**Table 2:** Morris-Lecar network synaptic parameter values and description.

| **Excitatory Synaptic Parameters** |  |
| --- | --- |
| $a_r = 3.0 \text{ (ms}^{-1}\text{)}$ | Rise rate constant |
| $a_d = 0.1 \text{ (ms}^{-1}\text{)}$ | Decay rate constant |
| $\bar{g} = 1.0 \text{ (mS/cm}^2\text{)}$ | Maximum conductance |
| $E_{ampa} = 0.0 \text{ (mV)}$ | Reversal potential (AMPA) |
| **Inhibitory Synaptic Parameters** |  |
| $a_r = 3.0 \text{ (ms}^{-1}\text{)}$ | Rise rate constant |
| $a_d = 0.1 \text{ (ms}^{-1}\text{)}$ | Decay rate constant |
| $\bar{g} = 1.0 \text{ (mS/cm}^2\text{)}$ | Maximum conductance |
| $E_{gaba} = -75.0 \text{ (mV)}$ | Reversal potential (GABA) |
| **Common Synaptic Parameters** |  |
| $T_{max} = 1.0$ | Maximum conductance scaling factor |
| $V_t = 2.0 \text{ (mV)}$ | Half-activation voltage for synaptic gating |
| $K_p = 5.0 \text{ (mV}^{-1}\text{)}$ | Slope factor for synaptic gating |

The constraint ensures that all connections between neurons are positive, aligning with the biological nature of neural connections. We remark that alternate scalings of the unitary synaptic conductances have been considered [38] before to elicit irregular activity in the form of chaotic initial network states [60, 61].

We constructed the ML network as illustrated in Fig. 2-A. In this architecture, we implemented a three-layer network in which interconnected ML neurons were in the hidden layer. The input and output layers were simple linear neural network neurons.

The input layer accepted the external stimulus at the time (***c***(*t*), e.g. a vector of pixel values) with a standard trainable linear transformation (***ω***_*in*_ and ***bias***_***in***_) to the hidden layer neurons, Eq. 35. This projection provided the external driving current, *I* (Eq. 26), to both excitatory/inhibitory neuron populations in the hidden layer. A scaling factor A = 40 was introduced to ensure that the external stimulus sufficiently activated the spiking neurons. This constant amplification was applied uniformly across all simulation tasks. Although some tasks might not have strictly required this factor, we maintained *A* = 40.0 throughout the experiments to guarantee a single, standardized network configuration and consistent input structure.

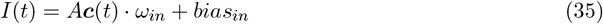

The conductance-based synaptic interactions between neurons were mediated by four sets of trainable weights: excitatory-to-excitatory (***ω***_*ee*_), inhibitory-to-excitatory (***ω***_*ei*_), excitatory-to-inhibitory (***ω***_*ie*_), and inhibitory-to-inhibitory (***ω***_*ii*_). Consistent with Eq. 34, all of these connections were bound during training using a *sigmoid* function transformation to follow Dale’s law.

Units are provided in parentheses where applicable.

This transformation, as shown in Eq. 36, maps the unbounded trainable *ω*_*raw*_ to the nonnegative synaptic weights used in the ML network.

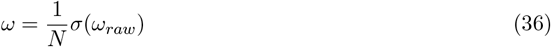

The *sigmoid* function *σ* ensured that all synaptic weights *ω* were positive, which was crucial for maintaining Dale’s law, the property that each neuron was excitatory or inhibitory but not both. By enforcing *ω ≥* 0, the nature of each neuron is solely based on the reversal potentials of the synapses and the presynaptic neuron’s identity: ***ω***_*ee*_ and ***ω***_*ie*_ are excitatory connections, while ***ω***_*ei*_ and ***ω***_*ii*_ are inhibitory connections. The 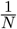, where *N* is the total number of neurons, scaled the weights to maintain the overall current input in the biologically plausible range.

The temporal evolution of each neuron’s variables was numerically integrated using the fourth-order Runge-Kutta (RK4) method, chosen for its balance of accuracy and computational efficiency in solving the differential equations, with a time step of *dt* = 0.1. A sample of temporal dynamics of randomly selected inhibitory/excitatory neurons during stimulus testing is shown in Fig. 2-B.

The output of the ML-SNN model was derived from the aggregation of synaptic gating variables (***s***(*t*)) of the hidden layer neurons. These variables represented the temporal integration of presynaptic activity and represented the post-synaptic potentials; in other words, they encapsulated the network’s internal state and its response to the input. The output layer applied this transformation using trainable output weights (***ω***_*out*_) and a bias (***bias***_*out*_), based on the aggregation of both excitatory and inhibitory synaptic gating variables, to produce the final network output signal over time, Eq. 37.

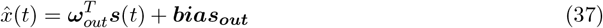

### Using the ML SNN for Feedforward and Recurrent Tasks

By utilizing the general ML-SNN model explained, we constructed a versatile foundation for exploring neural computation and learning. We investigated the model’s capabilities with two modes for different tasks. First, we utilized the standard feedforward model, which learned to map external stimuli presented at the input layer to the desired output. This model allowed us to evaluate its capacity for tasks such as pattern recognition and regression. Secondly, we explored an autonomous recurrent model by implementing a feedback loop that connected the network’s output to the input. Biologically, this feedback loop can be interpreted as higher order brain area projecting the current estimated dynamical state back into the SNN. This made the model capable of generating complex temporal dynamics internally by transforming it into a self-contained dynamical system.

In the first mode, the ML-SNN architecture was used for tasks regarding complex pattern recognition and regression from static inputs. These tasks involved the presentation of static images or numbers to the network’s input layer for a fixed duration *T*. The network processed this input over the simulation window, allowing the dynamics of the hidden layer’s ML neurons and their recurrent connections to integrate information over time. The output for classification tasks was made based on the accumulated activity of the output neurons. The predicted class was the one associated with the neuron with the highest average synaptic gating variable activity, *s*_*i*_(*t*) for *i* = 1, 2, … *j* where *j* = 10 for both MNIST and fashion-MNIST. For the regression task, the output was the average of the latter half of the output’s neuron activity, as shown in Fig. 3-B.

In the second mode, by closing the loop between the output and input layers, the network was capable of generating dynamics autonomously. In this condition, the network’s output at time *t* was fed back as the external stimulus for the subsequent time step. This feedback mechanism converted the SNN model from a stimulus-driven model into an autonomous system capable of generating complex temporal dynamics internally without requiring external input after an initialization phase. By training the network parameters appropriately, using a scheduled sampling strategy with linear decay controlling the interpolation between teacher-forced inputs and the model’s autonomous predictions [62, 63], this closed-loop operation allowed the model to construct dynamical systems autonomously.

In particular, the decoded output at the last timestep 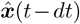 was fed back into the neurons as a current, using Eq 35, which becomes:

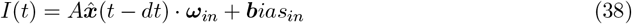

### Learning in Conductance Based Spiking Neural Networks

We used supervised learning to train the network’s parameters. This approach involved adjusting the weights based on a supervisory signal that corresponded to the desired output for a given input. The selection of this signal and the training process varied depending on the network’s task. In this section, we detail the supervised learning approach used to train the network.

### Supervisory Signals Used

The supervisory signal dictated how the network’s parameters should be adjusted to perform the desired task. In the classification tasks, we used a target-based supervisor to be active for the output neuron that corresponded to the correct class, while all other output neurons remained inactive. Given that our model utilized the temporal dynamics of the Morris-Lecar neuron, which evolved over a duration to make a prediction, the active signal was maintained at a constant value of 1 for the duration of the stimulus presentation, while inactive neurons remained at 0. By representing the classification targets as continuous signals over time, we effectively transformed the problem into regression, which aligned well with the model’s continuous-time dynamics and facilitated the use of gradient-based learning to optimize these dynamics. In the nonlinear regression task, we created a constant supervisory signal for the duration of simulation based on the expected true value *f* (*x*) = *x*^2^ based on an input to the network *x*.

In the dynamical systems tasks (the Lorenz and Van der Pol system), the supervisory signal was derived directly from the target dynamical system for dynamical system reconstruction. The target dynamical system was numerically simulated, 200 s for the Van der Pol and 1000 s for the Lorenz system, to produce a time series of state variables. The supervisory signal for this task was, therefore, the time series of the system’s state variables that the network was designed to replicate.

### Loss Function

The loss function, denoted as *L*, calculated the difference between the network’s output and the supervisory signal, as shown in Fig. 2-A, which guided the training process. We used different loss functions for different tasks.

For the classification tasks, we used the cross-entropy loss (Eq. 39) [40]. This loss function measures the difference between the network’s output probability distribution over classes and the target distribution created from the supervisory signal. A decrease in cross-entropy loss signifies improved alignment between these predicted and target distributions, leading the network to generate more precise classifications. The loss was computed using Pytorch’s embedded function, which is defined as:

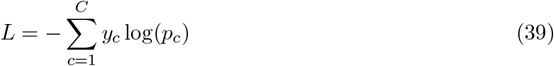

Where *C* is the number of task classes, and *y*_*c*_ represent a binary value indicating whether class *c* is the correct answer and is a standard one-hot vector where *k* is the index of the correct class:

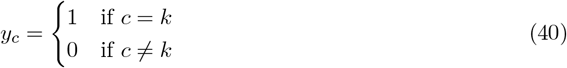

The *p*_*c*_ term is the predicted probability from the model’s output for the class *c*. To calculate it, first, we define the average readout (or logit) 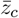 for each class *c* over time *t*:

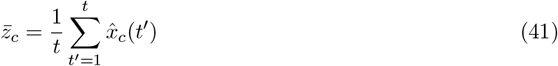

where 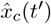 is the network readout for class *c* at time *t*^*′*^. The probability *p*_*c*_ is then calculated by applying the softmax function to these average logits:

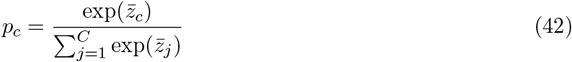

For dynamical system and regression tasks, the loss function utilized was the Mean Squared Error (MSE) loss, defined in 43. This function computed the mean of the squared differences between the actual target values provided by the supervisory signal and the network’s output:

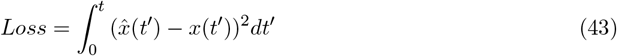

Where 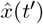 was the predicted signal from the model’s output and *x*(*t*^*′*^) was the actual signal coming from supervisory signals. For the MSE loss calculation, we used the embedded function torch.nn.MSELoss() in Pytorch’s library.

### Optimization

Using the results of the loss functions introduced earlier, the network’s weights (*ω*) were optimized using the AdamW optimizer [64] for both the recurrent and feedforward operating modes considered. AdamW is a variant of the Adam optimizer [65] that uses weight decay regularization, which helps to mitigate overfitting and enhance generalization. The optimizer equations are:

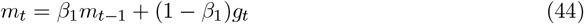

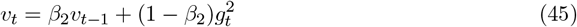

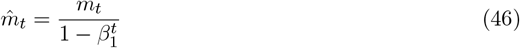

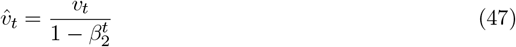

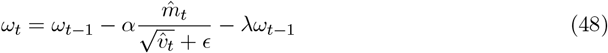

The parameters for AdamW are as follows: *g*_*t*_ is the gradient at time *t, m*_*t*_ and *v*_*t*_ are the first and second moment estimations of the gradient, *β*_1_ and *β*_2_ are the exponential decay rates for the moment estimates, 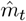 and 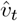 are the bias-corrected moment estimates, while *α* is the learning rate, a critical parameter governing the step size of weight updates. Finally, *ϵ* is a small constant to prevent division by zero in estimating the second moment, while *λ* is the weight decay coefficient, which controls the amount of regularization considered.

The AdamW optimizer updated the weights using a refined standard gradient descent approach by calculating adaptive learning rates for each parameter. This adaptation used the estimates of the first and second moments of the gradients. Subsequently, the bias-corrected moment estimates were used to update the weights, as shown in Eq. 48. This optimization procedure was used in both feedforward and recurrent operating modes.

Further, we used the Cosine Annealing Learning Rate scheduler, introduced in [66], which adjusted the learning rate *α* in Eq. 48 during training. The total number of training epochs was defined as half the period of a cosine oscillation. The scheduler enforces a smooth, monotonic decay of the learning rate, *α*, during the training. In our implementation, we utilized the PyTorch function, torch.optim.lr scheduler.CosineAnnealingLR, where the cycle duration *T*_*max*_ was set to the total number of epochs. The learning rate, *α*, is given by Eq. 49, where *α*_*initial*_ is the initial rate, *α*_*min*_ is the minimum rate (default 0.0), *T*_*max*_ is the total number of epochs, and *t* is the current epoch index:

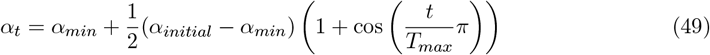

### Backpropagation Through Time

With an end-to-end differentiable SNN, we utilized gradient-based optimization to train the network’s weights. Particularly, we calculated the gradients of the loss function (*L*) with respect to the key trainable parameters of our SNN: the output weights and biases (***ω***_*out*_, ***bias***_*out*_), input weights and biases (***ω***_*in*_, ***bias***_*in*_) and recurrent hidden layer weights which includes four weight matrices: excitatory-to-excitatory (*ω*_*ee*_), inhibitory-to-excitatory (*ω*_*ei*_), excitatory-to-inhibitory (***ω***_*ie*_), and inhibitory-to-inhibitory (***ω***_*ii*_) that we summarized all into (***ω***). Mathematically, this involves computing:

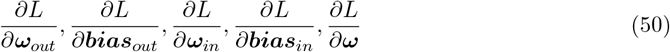

We employed Truncated Backpropagation Through Time (TBPTT) [41]. In TBPTT, the gradients were backpropagated only for a limited number of the most recent time steps, which we named the chunk size *T*_*BP T T*_. This truncation effectively limited the temporal dependencies and hence decreased the computational demands. By choosing an appropriate value for *T*_*BP T T*_ for each task, we reduced the memory usage and computational cost of training while the network learned.

For TBPTT, we utilized PyTorch’s library capabilities to calculate the gradient for each trainable parameter in each chunk size. Although the gradients could be calculated analytically, PyTorch’s library offers greater flexibility and simplified model development. These gradients were used by the AdamW optimizer, as detailed in the previous section, to update the network’s weights in a direction that minimized the loss function. This iterative process allowed our model to learn the complex mapping and temporal dynamics required for each specific task.

### Warm-up Phase

The warm-up phase was implemented at the start of every simulation run, in both training and testing, to ensure the network operated within a stable dynamic regime. As the ML neurons and their recurrent connections are initialized randomly, the network undergoes an initial, nonphysiological transient phase immediately following the start of the simulation. Allowing this transient to pass is essential because the ultimate goal is to evaluate the network’s performance, whether through loss calculation during training or output generation during testing, based on the network’s stable, characteristic spiking activity. By using a warm-up phase, we mitigate distortion of results caused by these unstable start-up dynamics.

The mechanism of the warm-up phase involves running the network forward in time while no input is presented, but with the crucial step of calculation intentionally disabled. Specifically, for training, the warm-up period is excluded from the backpropagation process: no loss is calculated, and no gradients are computed or stored for this initial duration, ensuring gradient updates (via TBPTT) are based on settled dynamics. For testing, the network’s output is not recorded or used for prediction until the warm-up time has elapsed. The duration of this phase was empirically determined and scaled based on the task complexity.

### Input and Training Specifics for Dynamical System Tasks

For the dynamical system reconstruction tasks, we added two procedures to maintain the stability of the system and to help the model learn the dynamics. First, to increase model stability, the target trajectories were separated into discrete chunks of duration *C*_*s*_ consistent with the Truncated BPTT chunk size *T*_*BP T T*_. Only the states associated with each chunk’s initial time step were given to the input layer of the model. After that, this input value was maintained throughout the chunk. By offering a piecewise constant input signal, this method helped stabilize the SNN’s recurrent dynamics by avoiding abrupt input fluctuations that could cause perturbation of the states of the neurons.

Second, a generalized teacher forcing technique [62, 63], also known as scheduled sampling, was added to the training procedure to help the model learn the dynamics. This was crucial for stabilizing the learning process when the network was transitioning from being driven by an external input signal to working solely based on its own output. Particularly, at each training step within an epoch, the input provided to the SNN, denoted as *I*_*input*_(*t*), was determined as a linear interpolation between the true target signal *x*(*t*) and the network’s own output prediction from the previous time step,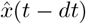. This interpolation was controlled by a coefficient *α*, so the input current was computed as:

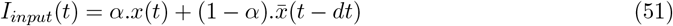

The interpolation coefficient, *α*, was changed over the course of training using a linear decay schedule. Its initial value was set to 1.0, to ensure the network input was determined only based on the supervisor. As training progresses, *α* linearly decreases toward a minimum value (*α* = 0.1). This scheduling method gradually increased the influence of the network’s own generated dynamics on the input and facilitated the transition to stable autonomous generation of the desired dynamics by the end of the training. During the testing phase/autonomous generation phase, *α* = 0 was used (no teacher forcing during testing).

### Programming Environment

All models were implemented in Jupyter Notebook with Python version 3 kernel using PyTorch for training the model. All differential equations were numerically integrated using the fourth-order Runge-Kutta (RK4) method with the time step *dt* specified for each task.

### Image Classification Tasks (MNIST & Fashion-MNIST)

For both classification tasks, we used a network consisting of an input layer with 784 neurons, corresponding to the 28×28 pixel images, a recurrent hidden layer of 1000 balanced ML neurons, and a linear output layer of 10 neurons, corresponding to the 10 classes of each dataset.

The input was prepared by normalizing the raw pixel intensities to the range [0, 1], and this constant input vector was presented to the hidden layer for *T* = 100 ms, simulated with a time step *dt* = 0.1 ms.

Models were trained for 5 epochs using the AdamW optimizer. The Cross-Entropy Loss was calculated on the time-averaged synaptic gating variables of the output neurons. The batch size was 200, the learning rate was set to 1 × 10^*−*3^, and the weight decay was 1 × 10^*−*4^. The gradients were calculated using TBPTT with a chunk size of *T*_*BP T T*_ = 100 time steps. A warm-up phase of 10 ms was also implemented at the beginning of each simulation to allow stabilization of the randomly initialized neuron population.

### Regression Task

We used a network consisting of an input layer of 1 neuron, a recurrent hidden layer of 500 balanced ML neurons (250 excitatory, 250 inhibitory), and a linear output layer of 1 neuron for this task.

A dataset of 5000 instances was generated with input values *x* sampled uniformly from the interval [*−*2.0, 2.0]. The dataset was split into 80% for training (4000 instances) and 20% for testing (1000 instances). Each input *x* was presented as a constant value for *T* = 500 ms.

The network was trained for 20 epochs to minimize the Mean Squared Error (MSE) loss, calculated between the target *x*^2^ and the model’s output. The final output value was computed by averaging the network’s output signal over the latter 250 ms of the simulation window. The AdamW optimizer was used with a batch size of 200, a learning rate of 1 × 10^*−*2^, and a weight decay of 1×10^*−*4^. The gradients were calculated using TBPTT with a chunk size of *T*_*BP T T*_ = 100 time steps. A warm-up phase of 20 ms was also implemented at the beginning of each simulation to allow stabilization of the randomly initialized neuron population.

### Dynamical System Reconstruction: Van der Pol System

For the Van der Pol autonomous reconstruction task we used a network of 2 input neurons; a hidden layer consisted of 500 ML neurons (250 excitatory, 250 inhibitory); and an output layer of 2 neurons.

The target supervisory trajectory was generated by numerically solving the Van der Pol equations, Eq. 21& 22 with *µ* = 1, using Python’s *solve ivp* function. The initial values were *x*(0) = 1.0 and *y*(0) = 0.0, and the system was simulated with a time step of *dt* = 5 × 10^*−*3^ for a total duration of 200s.

The network was trained for 20 epochs, with each epoch using a 10s segment of the total trajectory. The AdamW optimizer was used with an initial learning rate of 5 × 10^*−*3^ and a weight decay of 1 × 10^*−*4^. A chunk size of *T*_*BP T T*_ = 100 time steps used for TBPTT with a warm up duration of 100 ms. A cosine annealing learning rate scheduler was employed to adjust the learning rate during training. Also, The training employed the generalized teacher forcing strategy where the interpolation coefficient *α* was linearly decayed from an initial value of 1.0, full teacher forcing, to a minimum of 0.5 as training progressed.

### Dynamical System Reconstruction: Lorenz System

For the last task, we used 3 input neurons with a hidden layer consisting of 2000 balanced ML neurons along with a 3 neuron output layer.

As in the Van der Pol system, we numerically solved the Lorenz equations, Eq. 23- 25, with its three parameters set to 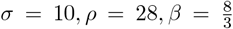 and the initial values were *x*(0) = 1.0, *y*(0) = 1.0, and *z*(0) = 1.0. The system was simulated with a timestep of *dt* = 5 × 10^*−*4^ for a total duration of 1000s.

The network was trained for 20 epochs, with each epoch using a 50s segment of the trajectory. The AdamW optimizer was used with a learning rate of 1 × 10^*−*3^ and a weight decay of 1 × 10^*−*4^. The chunk size of TBPTT for this task was *T*_*BP T T*_ = 200 time steps, with a warmup duration of 200 ms. The same Cosine Annealing and generalized teacher forcing techniques were used here as well.

### Supporting information

#### S1 Appendix. Gradient Derivation

Here, we derive the the gradient for the recurrent conductances explicitly as a set of differential equations for the loss function

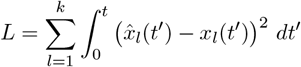

Where

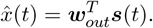

Then, we have the following:

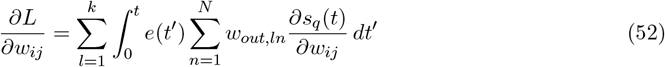

The partial derivative 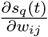 can be solved for as a differential equation as follows:

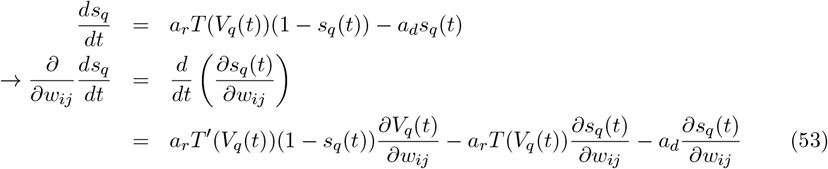

with a similar differential equation holding for 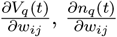:

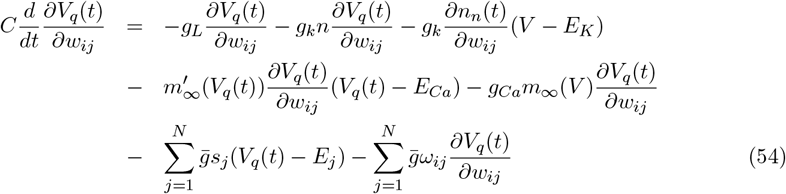

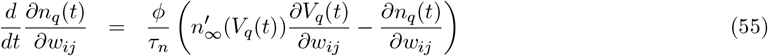

Collectively, equations (52)-(55) define a system of differential equations that can be used to explicitly determine the partial derivative 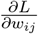. However, due to the complexity in implementing these *O*(*N* ^2^) differential equations during learning, a numerical gradient estimation scheme was instead used with the Adam with the gradient at time *t*.

### Supplementary Figures

**Supplementary Figure 1:**
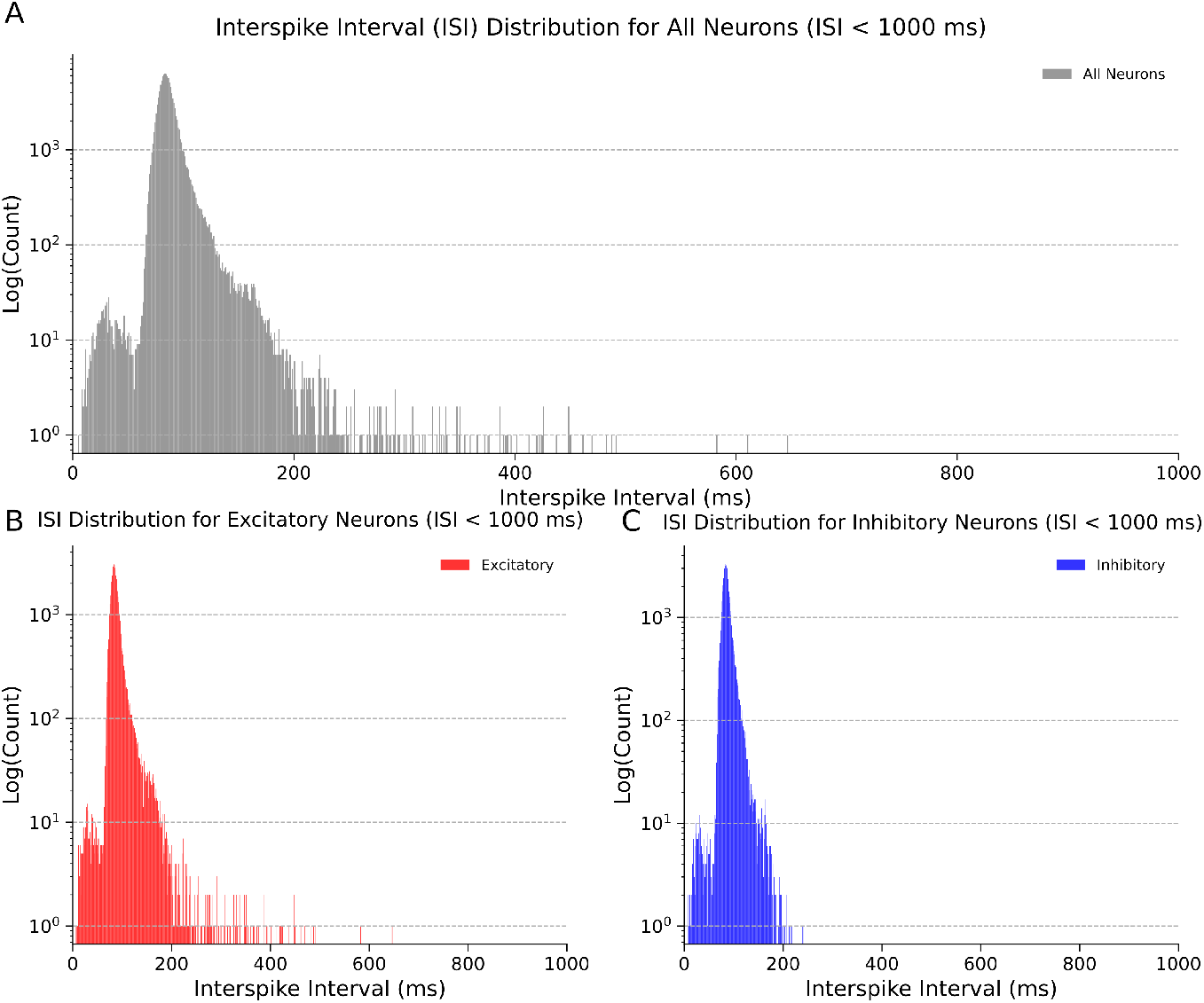
The interspike interval distributions for (A) all neurons in a untrained Morris-Lecar network, (B) the excitatory neurons, (C) the inhibitory neurons. The network was simulated for 1000 seconds and consisted of 100 balanced neurons, 50 inhibitory and 50 excitatory.

**Supplementary Figure 2:**
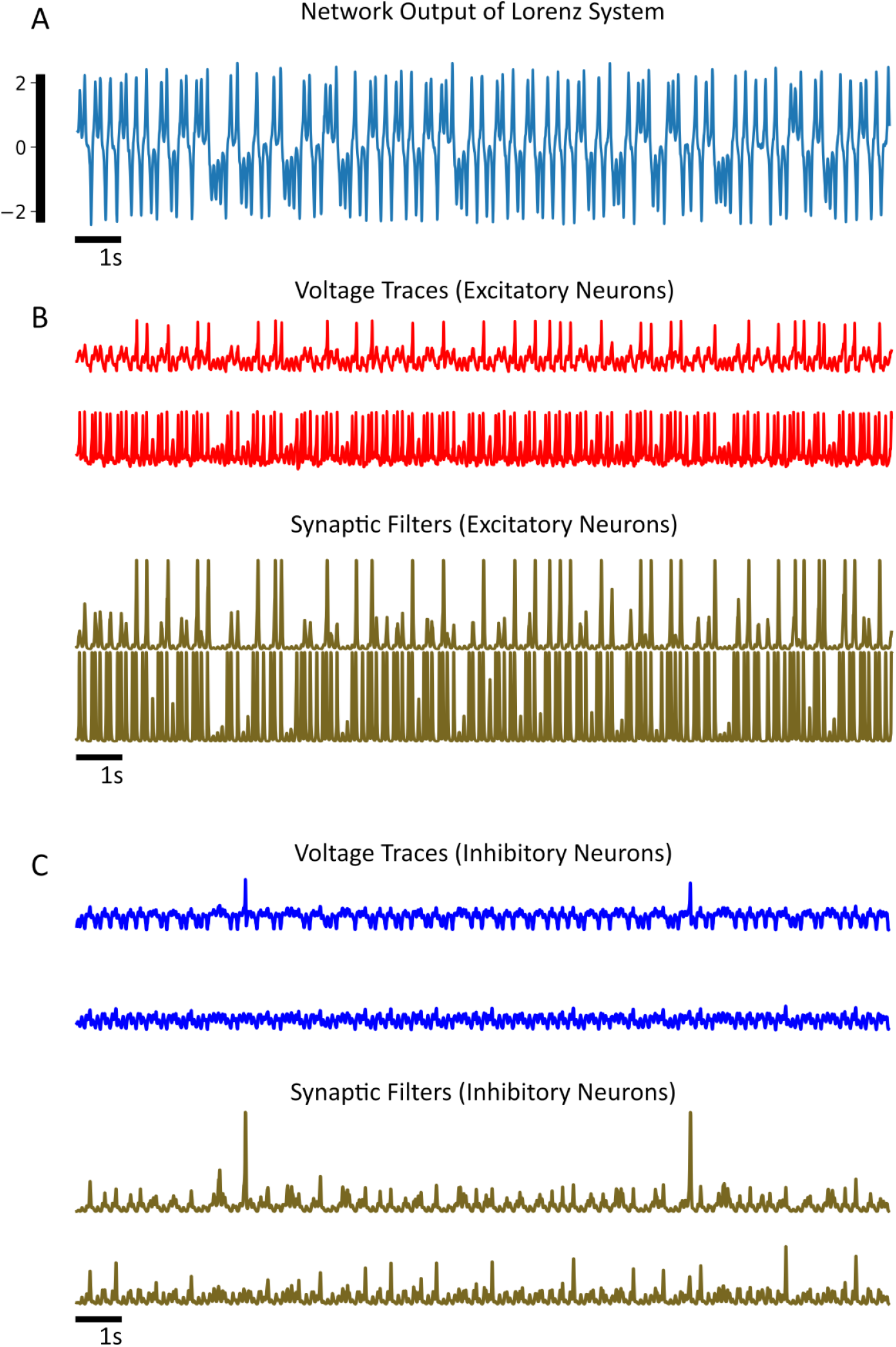
(A) The *x*(*t*) variable for the trained Lorenz system in Figure 4. (B) The voltage traces (top) and synaptic gating variables (bottom) for two excitatory neurons. The first (second) voltage trace corresponds to the first (second) synaptic gating variable. (B) The voltage traces (top) and synaptic gating variables (bottom) for two inhibitory neurons. The first (second) voltage trace corresponds to the first (second) synaptic gating variable

**Supplementary Figure 3:**
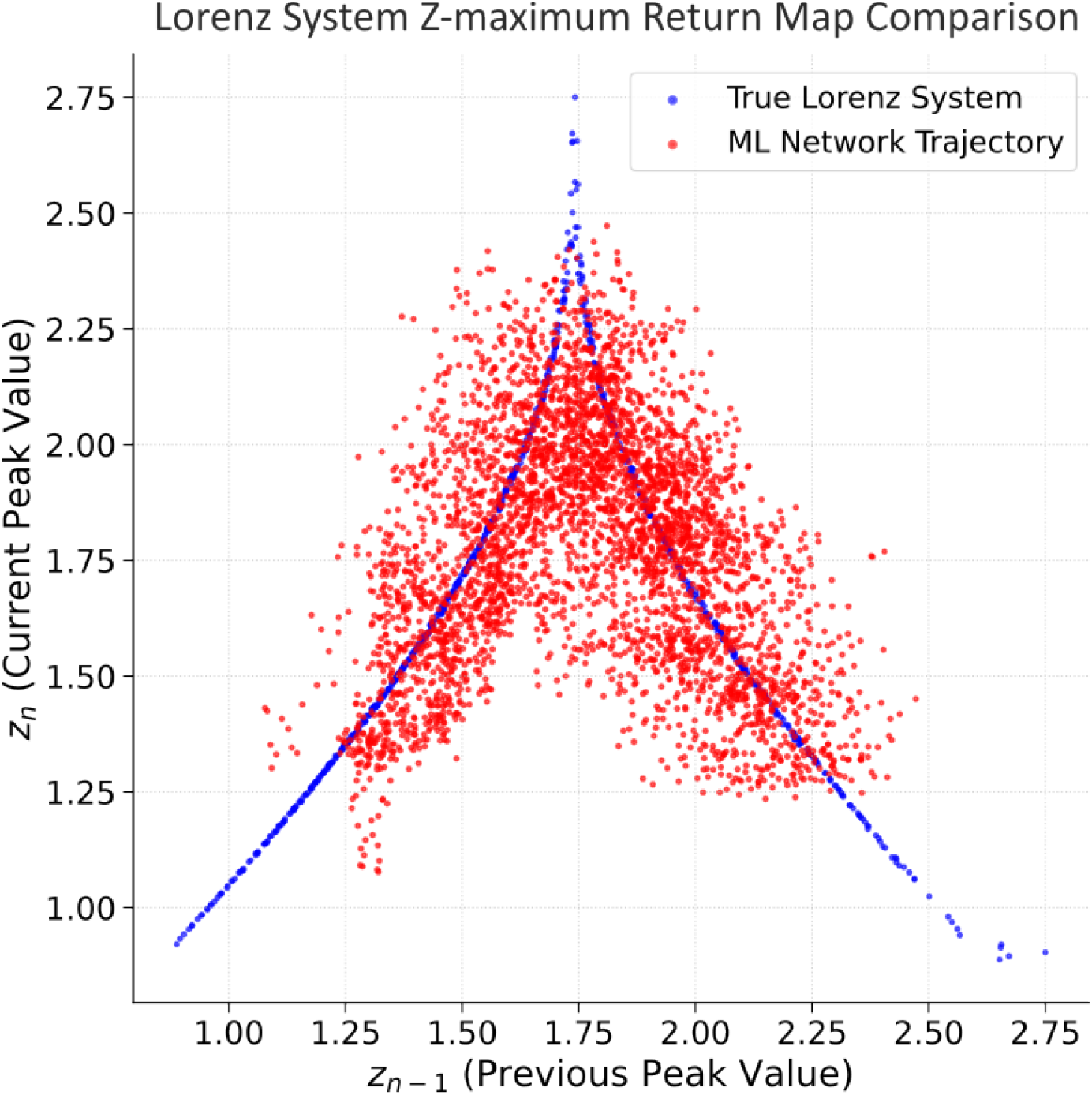
The return map for the Lorenz system (blue) *z*(*t*) variable, and the Morris-Lecar network (red). The return map was estimated with a simulation consisting of 500 seconds for both the Lorenz system and the ML network.

